# tHapMix: simulating tumour samples through haplotype mixtures

**DOI:** 10.1101/057414

**Authors:** Sergii Ivakhno, Camilla Colombo, Stephen Tanner, Philip Tedder, Stefano Berri, Anthony J. Cox

## Abstract

**Motivation:** Large-scale rearrangements and copy number changes combined with different modes of clonal evolution create extensive somatic genome diversity, making it difficult to develop versatile and scalable oriant calling tools and create well-calibrated benchmarks.

**Results:** We developed a new simulation framework tHapMix that enables the creation of tumour samples with different ploidy, purity and polyclonality features. It easily scales to simulation of hundreds of somatic genomes, while re-use of real read data preserves noise and biases present in sequencing platforms. We further demonstrate tHapMix utility by creating a simulated set of 140 somatic genomes and showing how it can be used in training and testing of somatic copy number variant calling tools.

**Availability and implementation:** tHapMix is distributed under an open source license and can be downloaded from https://github.com/Illumina/tHapMix.

**Supplementary information:** Supplementary data are available at *Bioinformatics* online.

## 1 Introduction

Recent advances in high-throughput sequencing of tumour genomes require robust and scalable tools for variant calling and annotation. While substantial progress has been made, polyploidy, normal sample contamination (aka purity) and polyclonality continue to confound interpretation of cancer genomes (Greaves et al., 2012). Changes in clonal composition are particularly difficult to estimate due to the lack of validated benchmarks. Failure to correctly account for tumour heterogeneity can lead to incorrect genome ploidy prediction that particularly impacts calling of copy number variants (CNVs). Several approaches have been developed for assessing tumour heterogeneity and for concomitant detection of somatic CNVs from the whole genome sequencing data (Oesper et al., 2013; Ha et al., 2014). The assessment and model training that these tools employ have been limited to mixing previously sequenced samples or simulating coverage distribution de-novo. The earlier approach requires a well-annotated truth set for each studied sample that is time-consuming to create, while the later introduces simplification that could distort underlying noise model. Tools specifically designed for creating SNVs, indels and structural variants from de-novo read simulation have also been introduced (Mu et al., 2015; Qin et al., 2015). However, they do not facilitate creation of large copy number variants that occur in most cancer samples; while genome sequences that they produce still require alignment and variant calling, which limits scalability when simulating hundreds of high-coverage whole-genome sequences.

Here we introduce a lightweight simulation engine of diverse tumour properties such as polyclonality, normal contamination and ploidy, tHapMix that scales up to hundreds of samples. It is based on haplotype mixing to create somatic genomes controlled for admixture that recapitulate properties of real sequencing data. To exemplify the utility of the tool, we provide a compendium of pre-processed data from 140 cancer genomes with diverse architectures and clonal composition and show how it can be used for automated training of CNV caller parameters.

## 2 Methods

### Outline

The tHapMix workflow comprises five distinct components designed to create haplotype-specific sequence data input and to generate a final set of somatic CNV and SNV variants on different branches of the cancer evolutionary tree. Clonal composition can be controlled through a number of parameters, including the number of clones, the percentage of heterogeneous variants, the structure of the clonal evolutionary tree (binary or single-level) and the percentage of normal contamination. tHapMix produces aligned reads in BAM format as the main output that could be easily plugged into CNV calling pipelines. Both whole-genome and chromosomal sequencing data can be simulated. Full details of HapMix workflow are available in Supplementary Material, Section 1.

### Deriving phased BAM files

Phased SNV and indel variants from the Platinum Genomes (PG) project (http://www.platinumgenomes.org) comprising an extended 17-member pedigree have been used to split aligned reads into separate haplotypes for each chromosome benchmark (Eberle *et al.*, 2016). PG samples NA12882 and NA12878 sequenced to a 200x depth on a HiSeq 2000 system were used as a source of aligned reads. This step needs to run only once for a particular sequencing platform and aligner.

### Truth files for simulation

CNVs are defined in BED format in which the location of the variants and distinct copy numbers for each haplotype are specified. tHapMix supports either ground truth files with known copy numbers per haplotype or user-defined input parameters for generating random truth files. Additionally, an optional file with small somatic variants in a VCF format can be specified; this will introduce SNVs on different branches of the clonal tree.

### Creating clonal truth files

A pool of variants contained in the ground truth files for CNVs (and optionally for SNVs) is subsampled to create clonal truth files of subclonal variants. All operations are consistent with the tree structure: i.e. variants shared by the entire population are located at the root node of the evolutionary tree and are present in all clones.

### Simulating and mixing clonal BAM files

tHapMix generates truth files for individual clones to sample reads from haplotype-split BAMs to produce heterogeneous variants in which coverage in a specific region is proportional to the pre-defined local copy number and the clonal composition. Each haplotype is sampled independently to simulate different allele frequencies. Purity of the sample (i.e. the percentage of tumour cells) and clonal abundances (i.e. the percentage of cells belonging to each clone) are supplied by the user to account for contamination with normal tissues and diverse distributions of the clonal subpopulations.

## 3 Results

We provide several examples illustrating tHapMix utility for software benchmarking, somatic CNV model training and in-silico analysis of tumour evolutionary mechanisms. In addition, Supplementary Material, Section 2.3 provides comparison with existing variant simulation tools.

### Large-scale simulation of tumourplocyclonality

We have used tHapMix to generate a total of 140 tumours with different modes of polyclonality. Truth sets representing genomes of different ploidy levels were created by comparison with karyotype data (where available, i.e. Newman et al., 2013) and by manual inspection of coverage, allele ratios and read mappings. The simulation strategy controlled for (1) number of clones, (2) percentage of polyclonal non-root variants, (3) preponderance of the major clone and (4) sample purity (Supplementary Material, Table 1 and 2). This protocol enabled the simulation of different evolutionary scenarios, ranging from monoclonal samples with no heterogeneity to highly diverse samples with multiple clones. For example, high relative abundance of the major clone represent models in which one clonal subpopulation is dominant in the sample as a result of positive selection. On the other hand, nearly equal abundances of two clonal subpopulations correspond to a model based on the expansion of divergent clones whose cohabitation in the tumour sample represents an evolutionary advantage.

### Software benchmarking

Large-scale datasets generated by tHapMix are well-suited for benchmarking CNV callers. We selected Canvas, which showed superior performance in a recent benchmark (Roller *et al.*, 2016), to assess how different modes of polyclonality influence variant calling (the assessment should generalize to other tools that use coverage and allele ratios to assign copy numbers). We find that overall abundance of heterogeneous variants most dramatically influences variant calling performance (p-value <0.05, linear regression t-test), followed by total number of clones and abundance of major clone (Supplementary Material, Section 2.1). This, in part, could be explained by the fact that extensive heterogeneity can influence assessment of normal contamination, leading to incorrect genome ploidy baseline and lower variant calling accuracy.

### Somatic model training

Modelling somatic CNV detection in the presence of polyclonality, normal contamination and aneuploidies remains a challenging task due to multi-modal hypothesis space and non-linear objective function. In such cases, model overfitting can easily occur over a limited training data, hampering reproducibility of accuracy metrics on new samples (Carter et al. 2012). Here we show how tHapMix simulations can be integrated into automated model training for somatic CNV calling in polyclonal and impure samples. Briefly, the workflow includes (1) reading Canvas somatic model configuration parameters file, (2) iteratively modifying parameters and assessing accuracy changes and (3) selecting the best preforming adjusted parameter set (Supplementary Material, Section 2.2). Assessment on an independent test data including 22 tumour samples of diverse genomic architectures and polyclonality features showed 12% accuracy improvement following such an optimization strategy (Supplementary Material, Figure 7).

### Summary

To conclude, tHapMix is a versatile and scalable workflow for simulation of diverse tumour characteristics, such as polyclonality purity and aneuploidy. It provides large-scale training and testing datasets to facilitate benchmarking and development of somatic variant calling tools, which are particularly hard to create for heterogeneous tumour samples.

## References

Carter,S. et al. (2012) Absolute quantification of somatic DNA alterations in human cancer. Nat Biotechnol., 30, 413–421.

Eberle,M. et al. (2016) A reference dataset of 5.4 million human variants validated by genetic inheritance from sequencing a three-generation 17-member pedigree. bioRxiv. doi: 10.1101/055541.

Ha,G. et al. (2014) TITAN: inference of copy number architectures in clonal cell populations from tumor whole-genome sequence data. Genome Res., 24, 1881–1893.

Greaves,M. (2012) Clonal evolution in cancer. Nature, 481, 306–313.

Mu,J.C. et al. (2015) VarSim: a high-fidelity simulation and validation framework for high-throughput genome sequencing with cancer applications. Bioinformatics, 31, 1469–1471.

Newman,S. (2013) The Relative Timing of Mutations in a Breast Cancer Genome. PLoS One, 8, e64991.

Oesper,L. et al. (2013) THetA: inferring intra-tumor heterogeneity from high-throughput DNA sequencing data. Genome Biology, doi:10.1186/gb-2013-14-7-r80.

Qin,M. et al. (2015) SCNVSim: somatic copy number variation and structure variation simulator. BMC Bioinformatics, doi:10.1186/s12859-015-0502-7.

Roller,E. et al. (2016) Canvas: versatile and scalable detection of copy number variants. Bioinformatics, 32, 2375–2377

